# Spatial analysis for highly multiplexed imaging data to identify tissue microenvironments

**DOI:** 10.1101/2021.08.16.456469

**Authors:** Ellis Patrick, Nicolas P. Canete, Sourish S. Iyengar, Andrew N. Harman, Greg T. Sutherland, Pengyi Yang

## Abstract

Highly multiplexed *in situ* imaging cytometry assays have made it possible to study the spatial organisation of numerous cell types simultaneously. We have addressed the challenge of quantifying complex multi-cellular relationships by proposing a statistical method which clusters local indicators of spatial association. Our approach successfully identifies distinct tissue architectures in datasets generated from three state-of-the-art high-parameter assays demonstrating its value in summarising the information-rich data generated from these technologies.

## Main

Understanding the interplay between different cells, cell sub-types and their immediate environment is critical for understanding the mechanisms by which cells organise themselves within tissues and their dysfunctions in the context of human diseases. Recent advances in high-parameter imaging cytometry technologies which include among others^1^ High-definition Spatial Transcriptomics^2^, seqFISH+^3^, CODEX^4^, t-CyCIF^5^, MIBI-TOF^6^ and IMC^7^ have fundamentally revolutionised our ability to observe these complex cellular relationships. Where standard immunohistochemistry protocols only allow the accurate visualisation of cells that can be uniquely characterised by two or three surface proteins, these cutting-edge high-parameter imaging cytometry technologies characterise cells with tens to tens of thousands of RNA or proteins enabling precise classification of cell sub-types and hence providing an unprecedented depiction of cellular heterogeneity in the spatial context in a tissue. The increase in size and high-dimensional nature of the images necessitates analytical tools that are more complex, computationally efficient, and require an increased need for interpretability than those currently used in standard imaging cytometry.

As low parameter (two or three) imaging cytometry protocols are well established, so too are the analytical techniques needed to analyse the data they produce. The two most widely used techniques for object-based analysis of cell localisation are nearest neighbour analysis and Poisson point process models^8^. Nearest neighbour analysis can be used to test if two cell types tend to be more closely located than would be expected by chance. In a k-nearest neighbour analysis, for each cell in cell type *c*_*1*_, the *k* closest cells are identified and then the number of another cell type *c*_*2*_ in this set are counted. These counts for each cell-region are then averaged and compared to the averages of cell-label permuted experiments to quantify significance. Alternatively, point process models have been used for decades in fields such as ecology, geology and astronomy to identify the localisation of objects. The K-function can be used to quantify and visualise a point process model. Over a range of radii, *r*, the K-function is a monotonic quantification of the average number of *c*_*2*_ cells within *r* distance of each *c*_*1*_ cell and adjusts for the expected number of cells in each cell region. Under certain assumptions statistical significance can be assessed for a Poisson point process model without the use of permutations.

There is a substantial need to develop analytical approaches for simultaneously identifying relationships between multiple cell types. Given the novelty of the technologies, analytical approaches for tackling this issue are in active development. Methods of note include Spatial Variance Component Analysis (SVCA) which addresses the problem by analysing images at the marker level, but amongst other influential effects, can quantify the changes in marker abundance explained by cell-cell interactions^9^. Alternatively, Goltsev *et al*. proposed an approach identifying 100 cliques of cell types from a nearest neighbour analysis showing that these captured histologically distinct regions of cell type composition^4^. Cellular neighbourhoods are identified by CytoMAP^10^ by clustering cell compositions of rasterised images defined by either scanning across the image or across cells and Giotto^11^ identifies spatial domains using a hidden Markov random field.

Here we propose a novel methodology implemented in R for identifying consistent spatial organisation of multiple cell types in an unsupervised way. Our method leverages established, and well-defined metrics to quantify point-process models. In short, our method clusters local indicators of spatial association (LISA) curves to enable the characterisation of multiple interactions between cell types in contrast to traditional pairwise analysis (**Figure 1**). LISA curves are a localised summary of a K-function from a Poisson point process model and have been used in varied contexts to identify landmines^12^ or clusters of forest fires and earthquakes^13^. In the following, we demonstrate that clustering these curves from multiple cell type cross-comparisons is a robust and stable method for providing a high-level summary of cell type localisation in high-parameter imaging cytometry data.

**Figure 1.**
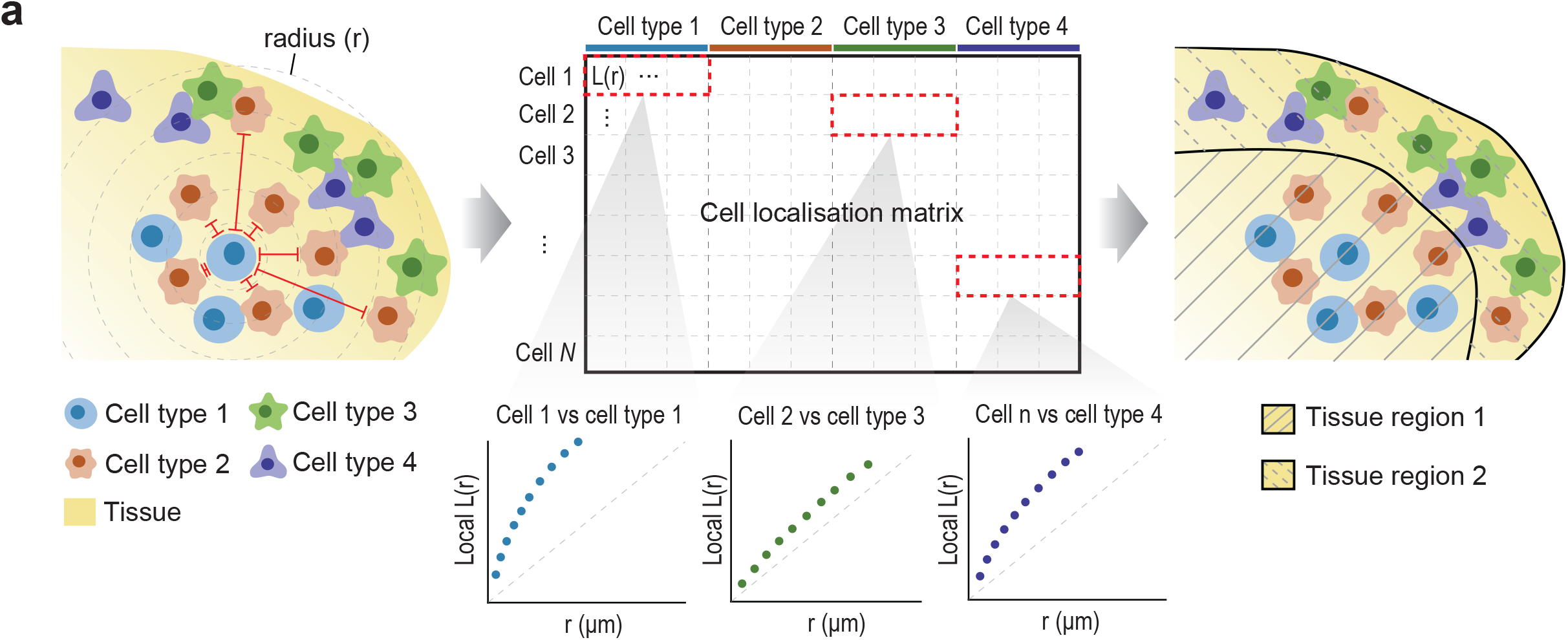
Clustering local indicators of spatial association. The workflow that lisaClust uses to identify regions of tissue with similar localisation patterns of cells contains multiple key steps. Firstly, cells are treated as objects and assigned coordinates in an x-y space. Second, distances between all cells are calculated and then, by modelling the cells as a multi-type Poisson point process, the distances are used to calculate local indicators of spatial association (lisa). These lisa curves are local L-functions summarising the spatial association between each cell and a specific cell type over a range of radii, *r*. The lisa curves are calculated for each cell and celltype and then clustered to assign a region label for each cell.

Our method, lisaClust, can identify distinct and well-defined tissue compartments in the spleen. The co-detection by indexing (CODEX) high-parameter imaging protocol was recently used to profile 26 immune cell types in murine spleens^4^. Applying the elbow method to estimate the number of clusters would suggest that the spatial relationships between the 26 cell types could be used to cluster the cells into three groups with distinct spatial ordering (**Supplementary Figure 1**). However, by clustering the cells into four groups we detected the four classic splenic compartments: red pulp, B cell follicles, PALS (periarteriolar lymphoid sheath), and marginal zone (**Figure 2a)**. The putative B cell follicle was enriched for B-cells and follicular dendritic cells, the putative PALS was enriched for CD4+ and CD8+ T cells, the putative marginal zone enriched for marginal zone macrophages and the putative red pulp region enriched for plasma cells (**Figure 2b**). The unsupervised identification of these tissue compartments was robust to the choice of clustering method with k-means clustering, hierarchical clustering, Clustering Large Applications (CLARA)^14^ and Self-Organising Maps (SOM)^15^ all identifying similar regions (**Supplementary Figure 2**).

**Figure 2.**
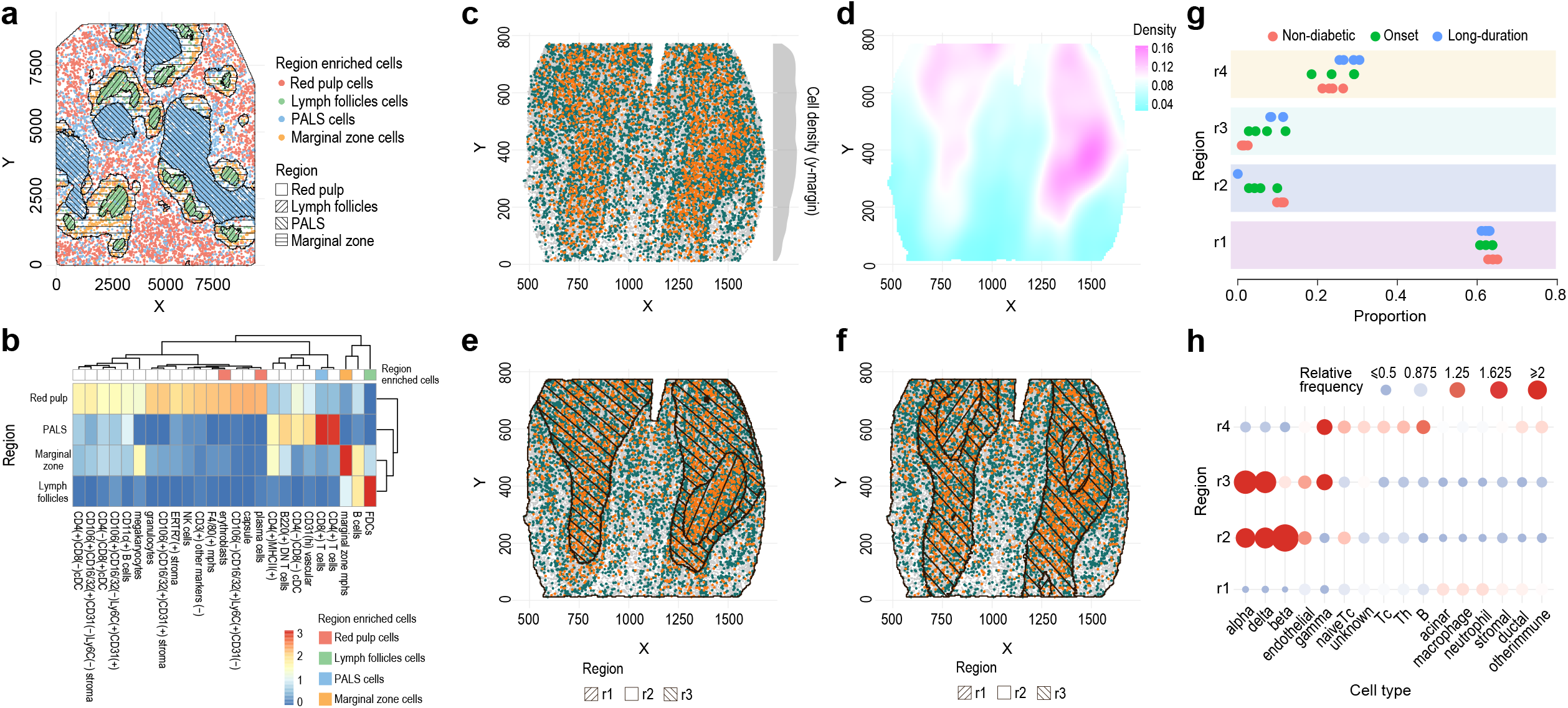
lisaClust is an effective approach for quantifying high-dimensional spatial ordering in high-parameter *in situ* imaging cytometry data. **a)** In a sample of spleen tissue imaged with CODEX, lisaClust is used to cluster 81693 cells into four distinct regions highlighted by different hashing patterns. The four regions are representative of the red pulp, lymph follicles, PALS and marginal zone. Specific cell types that were enriched in the red pulp (red), lymph follicles (green), PALS (blue) and marginal zone (yellow) are plotted. **b)** a heatmap is used to quantify the enrichment of the 26 cell types in each region highlighted in a). **c)** HDST is used to profile 69143 cells with 63 cell types in a murine olfactory bulb (OB). Cells are plotted in x-y coordinates (gray) with *OBNBL1+* neurons (green) and *OBINH1+* neurons (orange) highlighted. **d)** density of the OB cells with high density in pink and low density in blue. **e)** Estimates using a homogenous local L-function capture OB regions with different cell density and **f)** regions estimated from an inhomogenous local L-function outline the OB granule cell layer and the subependymal zone. **g)** lisaClust is used to identify four regions contained within 845 IMC images of pancreatic isletsand a heatmap summarises the enrichment of 16 cell types in each region. **h)** The proportion of cells from region r2, which contain alpha cells, delta cells and gamma cells but few beta cells, is higher in images from long duration type 1 diabetes individuals than non-diabetics.

The density of cells across a tissue can influence the types of regions identified by lisaClust. Depending on the scenario, identifying spatial domains which contain complex cellular interactions might be preferred to identifying regions which are representative of dense tissue architecture. This contrast can be observed in a High-definition Spatial Transcriptomics (HDST) dataset which profiled the spatial organisation of 63 cell types in the murine olfactory bulb (**Figure 2c**). When lisaClust was used to identify three distinct regions (**Figure 2d**), these regions primarily reflected the density of cells in the tissue (**Figure 2e)** with one region capturing the densest portion that is the inner granule cell layer (**Figure 2d, Supplementary Figure 3a**). However, statistical models which account for strong spatial structure are well defined. By modelling the spatial organisation of cells as an inhomogeneous Poisson point process and calculating the corresponding LISA curves, lisaClust was instead able to demarcate the whole granule cell layer and the subependymal zone (**Figure 2f, Supplementary Figure 3b**). This illustrates the importance of the operator being cognisant about whether identified regions will be influenced by cell density and that lisaClust is flexible to this decision.

Finally, lisaClust can identify spatial ordering of cell types that are associated with a measured outcome across samples. Imaging Mass Cytometry (IMC) has been used to profile 16 cell types in pancreatic islets from individuals with recent onset type 1 diabetes, long-term type 1 diabetes and non-diabetics^4^. After applying lisaClust to identify four distinct regions, the overall area of these regions was used to quantify the amount or level of specific complex cell type interactions in each sample. Two of the regions were representative of islets and were highly enriched for alpha cells, delta cells and gamma cells while one region also contained beta cells and another no beta cells at all (**Figure 2g**). Beta cells produce insulin and so, unsurprisingly, the regions that contain the beta cells were present in non-diabetics but not in long-term type 1 diabetics (**Figure 2h**). While simpler analysis strategies might have have identified that the proportion of beta cells was different between diabetics and non-diabetics, they would not have placed this difference into the broader context of their loss from the islet. This approach demonstrates an alternative to multiple pairwise localisation tests and is a viable step towards a multivariate analysis procedure.

Overall, clustering local indicators of spatial association is an effective approach for quantifying high-dimensional spatial ordering in highly multiplexed *in situ* imaging cytometry data. However, it does have limitations as three parameters need to be chosen by the operator. In general, like the elbow method which we used, many methods exist for optimising the number of clusters to choose when clustering. We have not evaluated these here given a lack of ground truth and that, in our opinion, the number of clusters chosen should be contextual. The range of radii over which to calculate the curves also needs to be decided by the operator but clustering curves, instead of a single distance, appears to lessen the impact of this choice^16^ (Supplementary Figure 4). Further, the choice of whether to use an inhomogeneous Poisson point process model, or not, can alter the types of regions identified (**Figure 2e, Figure 2f**) and should be selected purposefully.

We have illustrated that lisaClust can be applied to identify tissue compartments and prioritise distinct cellular microenvironments in multiple imaging assays that profile cellular composition at a single-cell resolution. The method is available as an open-source R package (https://bioconductor.org/packages/release/bioc/html/lisaClust.html) and as an online shiny application https://shiny.maths.usyd.edu.au/lisaClust/.

## Acknowledgements

We thank the scientific community for proactively making their data and code publicly available. This work has been partly supported by The University of Sydney and an Australian Research Council Discovery Early Career Researcher Award (DE200100944) funded by the Australian Government.

## Contributions

All authors contributed to the interpretation of results and writing of the manuscript. E.P and N.P.C conceived the project with E.P and S.S.I performing the main analysis and E.P and P.Y generating the figures.

## Methods

### lisaClust

We propose Local Indicators of Spatial Association Clustering (lisaClust) as an algorithm for identifying distinct tissue microenvironments. The identified tissue microenvironments are regions of tissue that are enriched for combinations of certain cell types. Practically, lisaClust is a dimension reduction strategy which facilitates identifying and summarising complex spatial organisation of multiple cell types.

#### Local indicators of spatial association

lisaClust is an object-based spatial analysis technique where each cell is represented as a labelled point in a two-dimensional plane. As such, it is assumed that any input data has undergone single-cell segmentation and cell type classification. The core of the approach is clustering localised summaries of multitype Poisson point process models which quantify the spatial localisation of pairs of cell types.

By modelling cells from each image as a multi-type Poisson point process, the spatial localisation between two cell types can be quantified with a K-function^17^,

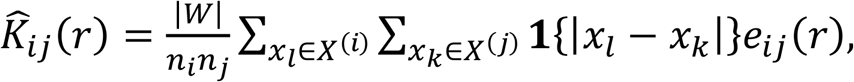

where 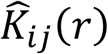 summarises the degree of localisation of cell type *j* with cell type *i* within radius *r, n*_*i*_ and *n*_*j*_ are the number of cells of type *i* and *j, x*_*m*_ are the x-y coordinates of cell *m*, |*W*| is the image area and *e*_*ij*_(*r*) is an edge correcting factor. The K-function can be interpreted as the average number of cells of type *j* within a distance r away from each cell of type *i*.

The L-function is a variance-stabilised K-function given by

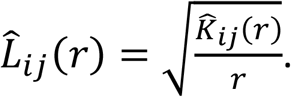

The expected value of *L*_*ij*_(*r*) is *r* and so the centered L-function 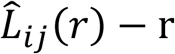 will have constant variance and a mean of zero.

To calculate local indicators of spatial association, the contribution of each individual cell to the multi-type K-functions of cell type j is quantified as follows:,

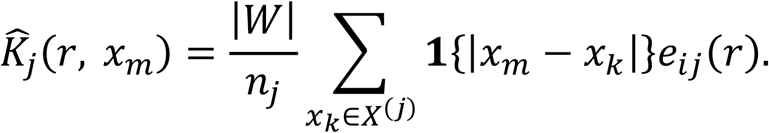

and then clustered on 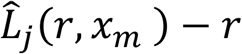. This measure is greater than zero if there are more cells of type *j* near cell *m* than expected and less than zero if there are less than expected.

#### Weights for tissue inhomogeneity

Tissue typically has non-uniform structure which means that assuming the point process has a homogeneous intensity over the range of the image is inappropriate. To better summarise the degree of localisation in images with strong structure, the local contributions of cells to the inhomogeneous K-function is also calculated

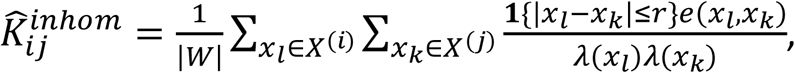

where *λ*(*x*_*m*_) is the expected density of cells at cell *x*_*m*_. *λ*(*x*_*m*_) is estimated using a Gaussian kernel with standard deviation 20.

#### Clustering

For the key results in this manuscript, k-means clustering is used to group the cells by their local indicators of spatial association using the *kmeans* function in the *stats* package in R. A fast implementation of hierarchical clustering from the *fastcluster* package^18^ is used for hierarchical clustering, Self-Organising Maps (SOM) are generated using the *kohonen* package and Clustering Large applications (CLARA) is implemented using the *cluster* package..

### Datasets

lisaClust is applied to three highly multiplexed *in situ* imaging cytometry datasets. Each of these datasets was generated with a different technology.

#### Co-detection by indexing (CODEX)

CODEX (CO-Detection by indEXing) is a highly multiplexed cytometric imaging approach that uses antibodies conjugated to oligonucleotide sequences^4^. After staining a tissue sample with an antibody panel, dye labelled reporters with complementary sequences can be iteratively imaged and removed to quantify protein abundance on cells. Segmented data of a “BALBc-1” spleen sample taken from a normal BALB/c mouse was downloaded from https://data.mendeley.com/datasets/zjnpwh8m5b/1. A 30-antibody panel had been used to image 81693 cells and these were annotated into 26 cell types by the authors.

#### High-definition Spatial Transcriptomics (HDST)

HDST is an assay which captures RNA from tissue sections on a dense, spatially-barcoded bead array^2^. The beads can measure the expression of thousands of RNA at 2 μm resolution making it possible to measure RNA expression at single cell resolution. The segmented data was downloaded from https://static-content.springer.com/esm/art%3A10.1038%2Fs41592-019-0548-y/MediaObjects/41592_2019_548_MOESM7_ESM.xlsx of a main olfactory bulb from a C57BL/6J mouse. This single image contained 69143 cells, and these were annotated into 63 cell types by the authors.

#### Imaging Mass Cytometry (IMC)

IMC is a highly multiplexed cytometric imaging approach that uses antibodies conjugated to metal isotopes^7^. After staining a tissue section with the metal-tagged antibodies, the tissue is ablated using a laser with a 1 μm spot size, the plumes of tissue matter are aerosolised, atomised, and ionised, and directed through a time-of-flight mass spectrometer for isotope quantification. The segmented data of 845 images of pancreatic islets from four non-diabetic human donors, four donors with recent onset type 1 diabetes and four donors with long duration type 1 diabetes were downloaded from https://data.mendeley.com/datasets/cydmwsfztj/1. A 35-antibody panel had been used to classify 1776974 cells into 16 distinct cell types by the authors.

## Code availability

The code for generating the figures in this manuscript is available on https://github.com/ellispatrick/lisaClustPaper

