## Supplementary File 1 for "Spatial analysis for highly multiplexed imaging data to identify tissue microenvironments"

**Supplementary Figure 1 – The optimal number of clusters to choose for CODEX dataset**

In a sample of spleen tissue imaged with CODEX, lisaClust was used to cluster 81693 cells into distinct regions using k-means clustering. The R package, *factoextra*, was used to estimate the optimal number of clusters. On the y-axis is the Total Within Sum of Squares (WSS) and on the x-axis the number of clusters used. The point at which the curve starts to flatten (the elbow) suggests the ideal number of clusters to use would be three.

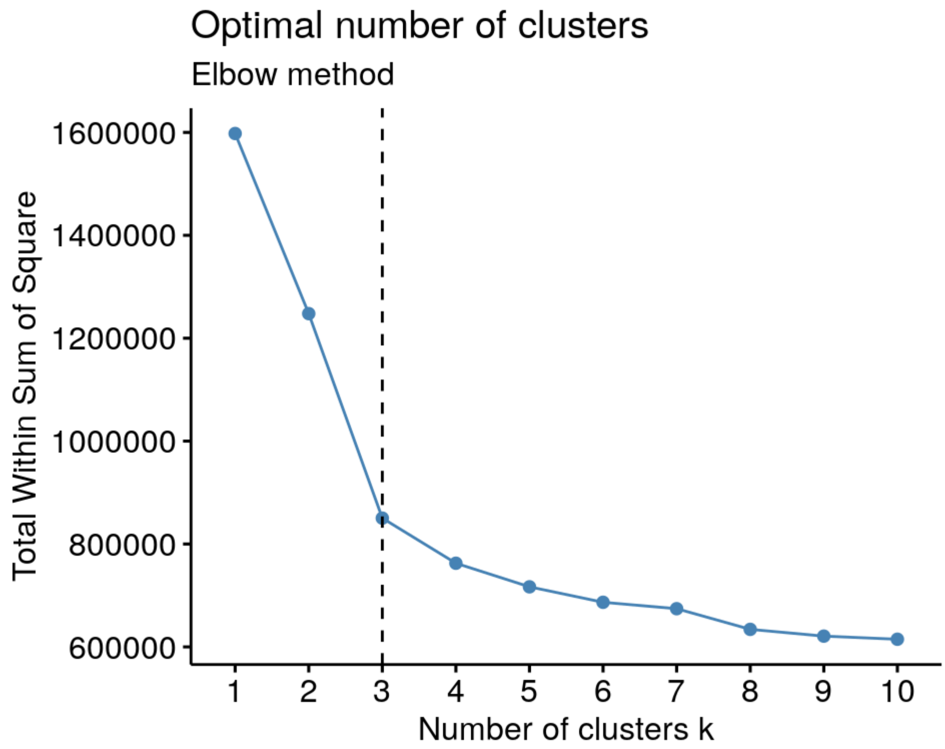

**Supplementary Figure 2 – Identification of the tissue compartments in the spleen is robust to clustering approach**

In a sample of spleen tissue imaged with CODEX, lisaClust was used with **a)** k-means clustering, **b)** Clustering Large Applications (CLARA), **c)** hierarchical clustering and **d)** Self-organising map (SOM) to cluster 81693 cells into four distinct regions highlighted by different hashing patterns. All four clustering approaches identified four similar regions. The four regions are representative of the red pulp, lymph follicles, PALS and marginal zone. Specific cell types that were enriched in the red pulp (red), lymph follicles (green), PALS (blue) and marginal zone (yellow) are plotted.

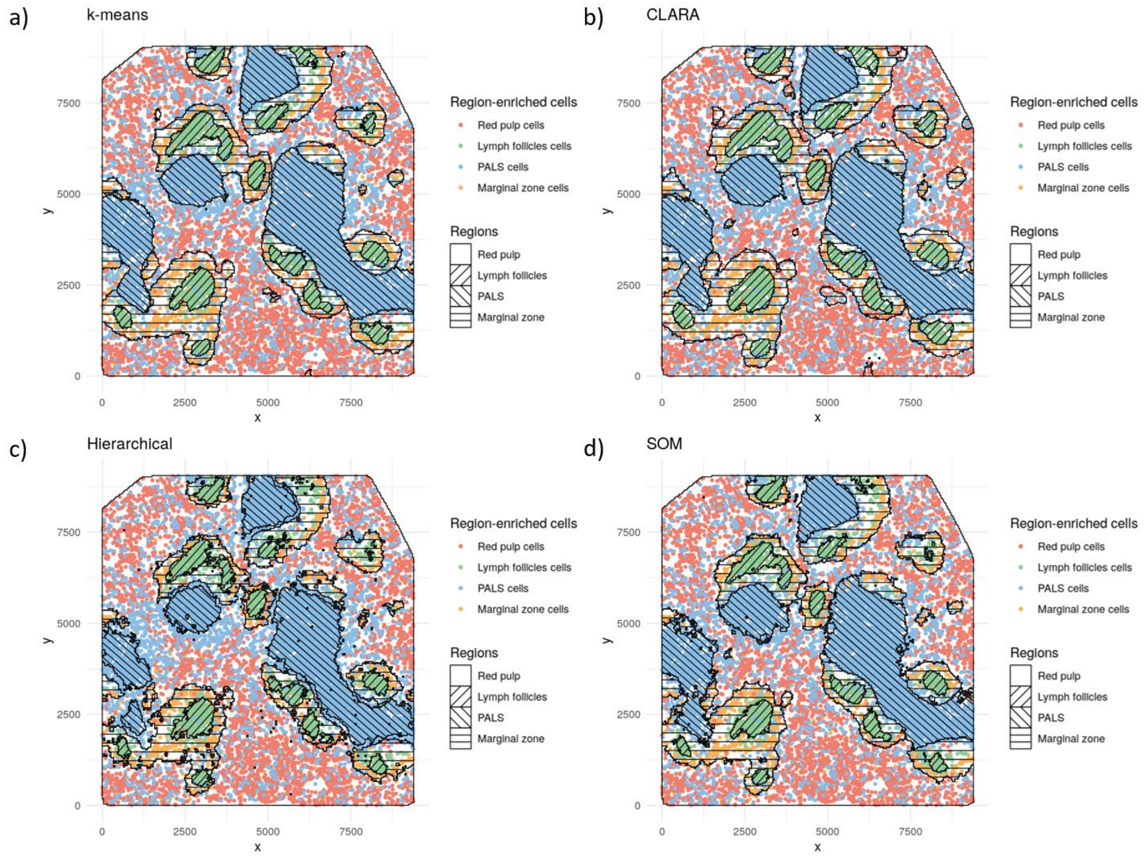

**Supplementary Figure 3 – Relative enrichment of cell types in regions identified in murine olfactory bulb**

High-definition Spatial Transcriptomics was used to profile 69143 cells with 63 cell types in a murine olfactory bulb (OB). lisaClust was used to identify three regions of spatial ordering using **a)** homogenous local L-functions and **b)** inhomogenous local L-functions. Heatmaps are used to visualise the relative enrichment of each of the 63 cell-types in each region with red meaning there are more cells of that type than expected and blue that there are less.

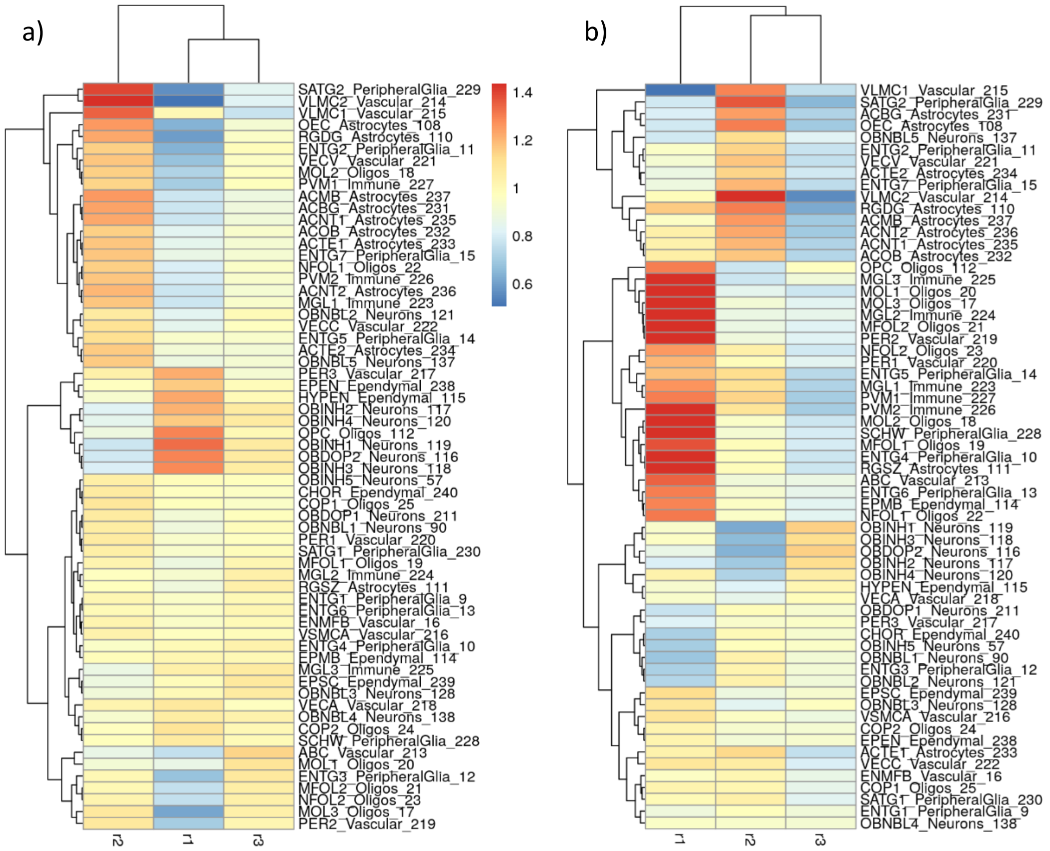

**Supplementary Figure 4 – Identification of the tissue compartments in the spleen is robust to clustering approach**

High-definition Spatial Transcriptomics was used to profile 69143 cells with 63 cell types in a murine olfactory bulb (OB). lisaClust was used to identify three regions of spatial ordering using either the combination of radii r = 9, 16, 25, 36, 49, 64, 81 and 100 as input or using each of these individually. For the plot using the combination of the radii, cells are plotted in x-y coordinates (gray) with *OBNBL1+* neurons (green) and *OBINH1+* neurons (orange) with the identified regions highlighted with hatching. In the remaining plots each of the cells are coloured by the regions that were identified using a single radius, with the combination regions highlighted in hatching. Using the combination appears to identify similar regions to r = 64, highlighting the OB granule cell layer and the subependymal zone.

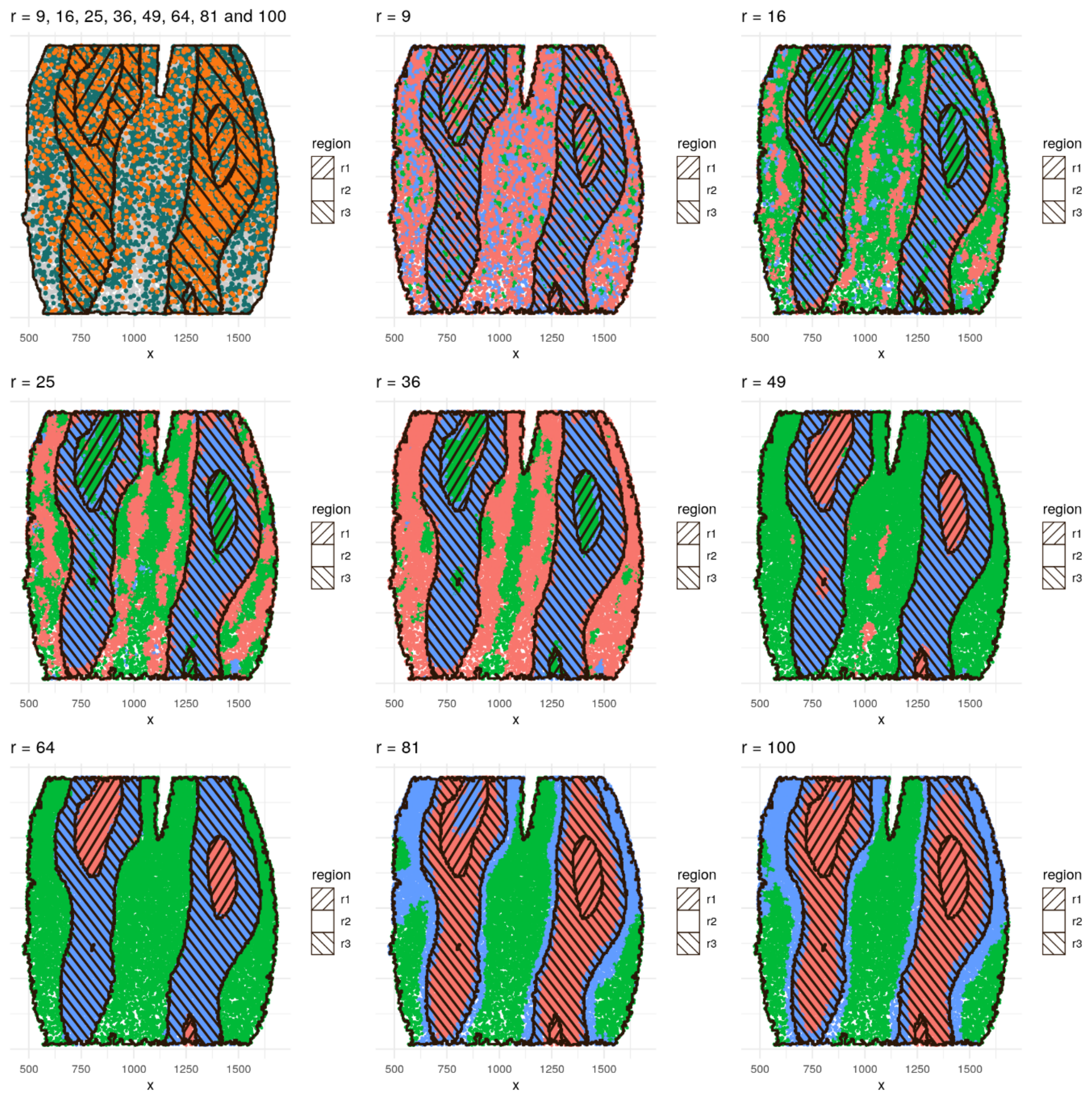
